# Identifying Cancer-Relevant Mutations in the DLC START Domain using Evolutionary and Structure-Function Analyses

**DOI:** 10.1101/2020.09.20.305201

**Authors:** Ashton S. Holub, Renee A. Bouley, Ruben C. Petreaca, Aman Y. Husbands

## Abstract

Rho GTPase signaling promotes proliferation, invasion, and metastasis in a broad spectrum of cancers. Rho GTPase activity is regulated by the Deleted in Liver Cancer (DLC) family of bonafide tumor suppressors which directly inactivate Rho GTPases by stimulating GTP hydrolysis. In addition to a RhoGAP domain, DLC proteins contain a StAR-related lipid transfer (START) domain. START domains in other organisms bind hydrophobic small molecules and can regulate interacting partners or co-occurring domains through a variety of mechanisms. In the case of DLC proteins, their START domain appears to contribute to tumor suppressive activity. However, the nature of this START-directed mechanism, as well as the identities of relevant functional residues, remain virtually unknown. Using the Catalogue of Somatic Mutations in Cancer (COSMIC) dataset, and evolutionary and structure-function analyses, we identify several conserved residues likely to be required for START-directed regulation of DLC-1 and DLC-2 tumor suppressive capability. This pan-cancer analysis shows that conserved residues of both START domains are highly-overrepresented in cancer cells from a wide range tissues. Interestingly, in DLC-1 and DLC-2, three of these residues form multiple interactions at the tertiary structural level. Further, mutation of any of these residues is predicted to disrupt interactions and thus destabilize the START domain. As such, these mutations would not have emerged from traditional hotspot scans of COSMIC. We propose that evolutionary and structure-function analyses are an underutilized strategy which could be used to unmask cancer-relevant mutations within COSMIC. Our data also suggest DLC-1 and DLC-2 as high-priority candidates for development of novel therapeutics targeting their START domain.

**Simple Summary:** Deleted in Liver Cancer (DLC) proteins are tumor suppressors that contain a StAR-related lipid transfer (START) domain. Little is known about the DLC START domain including the residues that mediate its activation. Using the Catalogue of Somatic Mutations in Cancer (COSMIC) dataset, evolutionary, and structure-function analyses, we identify key functional residues of DLC START domains. Mutations in these residues are significantly overrepresented in numerous cancers in multiple tissue types. In other contexts, START domains bind hydrophobic small molecules and stimulate regulatory outputs of interacting partners or co-occurring domains. The identification of functional residues in the DLC START domain may thus have implications for targeted manipulation of DLC tumor suppressive activity. Critically, these residues would not have been identified using traditional queries of COSMIC. Thus, we propose tandem evolutionary and structure-function approaches are an underutilized strategy to unmask cancer-relevant mutations within COSMIC.

## 1. Introduction

Rho GTPases are a subfamily of G proteins involved in signal transduction. These proteins regulate multiple pathways, including the actin cytoskeleton, cell polarity, cell cycle progression, and microtubule dynamics [1]. Signaling is activated by GTP binding at their GTPase domain and inactivated by its hydrolysis into GDP. This balance is mediated by the opposing activities of Rho guanine nucleotide exchange factors (RhoGEFs) and Rho GTPase activating proteins (RhoGAPs), which promote GTP replacement and GTP hydrolysis, respectively (**Figure 1**). Upregulated signaling of Rho GTPases such as RhoA, Cdc42, and Rac, or their respective RhoGEFs, leads to tumor proliferation, migration, invasion, and metastasis in a number of cancers [2–4].

**Figure 1.**
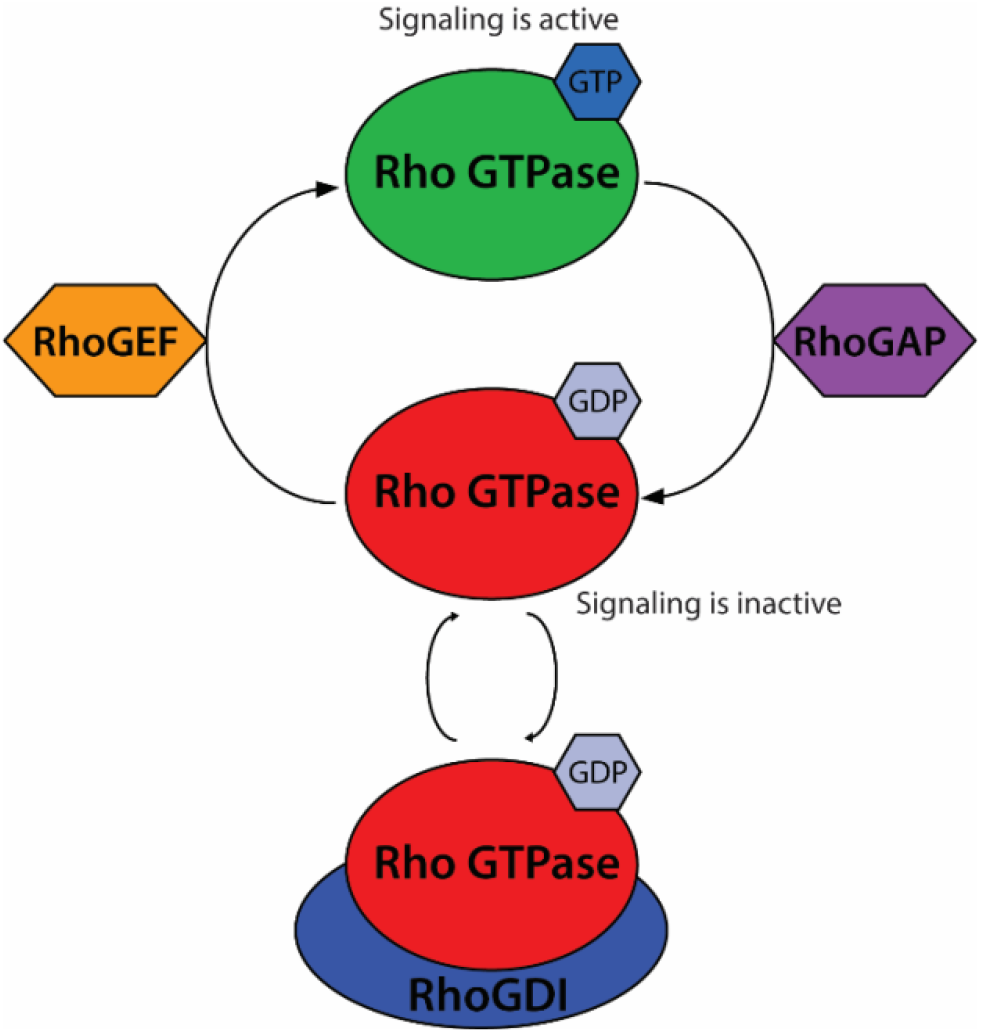
The Rho GTPase signaling module. Rho GTPases (red) are active while bound to GTP. RhoGAPs (purple) stimulate hydrolysis of bound GTP, switching Rho GTPases an inactive GDP-bound state. RhoGDIs (blue) bind GDP-bound Rho GTPases and sequester them in the cytosol. Finally, RhoGEFs (orange) exchange bound GDP for GTP, reactivating Rho GTPase signaling.

Rho GTPases have a globular structure with a limited druggable surface, making them particularly challenging drug targets [5–7]. Efforts to mitigate the effects of Rho GTPases in cancer have therefore focused primarily on disrupting aspects of Rho GTPase signaling (reviewed in [7]). These include studies of RhoGAPs, which have specific consistent inhibitory effects on Rho GTPase signaling and are considered particularly attractive components to target within the Rho GTPase signaling module [1,3,5,7–14] (**Figure 1**). The Deleted in Liver Cancer 1 (DLC-1 or STARD12), DLC-2 (STARD13), and DLC-3 (STARD8) RhoGAPs are frequently downregulated in cancer and associated with poor prognosis [12,15]. The Cancer Genome Atlas (TCGA) data indicate that, of the three DLC genes, DLC-1 is usually the most dramatically downregulated, followed by DLC-2 [12]. However, unlike DLC-2, DLC-1 is embryonic lethal when knocked out in mouse models [16,17]. A recent pan-cancer analysis also showed that DLC-1 missense mutations are common in cancer cells [15]. This suggests a larger role for DLC-1 in control of disease and development. DLC-3, on the other hand, does not show a correlation between copy number loss and cancer progression in the TCGA dataset [12]. Despite making differential contributions to cancer progression, all three DLC proteins are considered to be bonafide tumor suppressors that inhibit cancer cell growth [16,18–24].

Adjacent to the DLC RhoGAP domain is a StAR-related lipid transfer (START) domain. START domains adopt a helix-grip fold structure, forming deep hydrophobic binding pockets that undergo conformational changes upon binding of lipophilic ligands [25–27]. START-containing proteins are distributed throughout Eubacteria and Eukaryota, and are important regulators of numerous biological processes. For instance, in humans, START-containing proteins are associated with numerous human diseases, including metabolic, oncogenic, and autoimmune disorders [28–30]. Structurally, these proteins can be divided into two classes: minimal START proteins and START-containing multidomain proteins (SMPs), which possess a wide variety of additional functional domains [31]. START domains employ a range of complex regulatory mechanisms [19, 20, 32–34]. Of particular interest, START domains in both minimal proteins and SMPs are able to stimulate regulatory outputs of interacting partners or co-occurring domains in a ligand-dependent manner [32, 34]. In the context of DLC proteins, this would be predicted to manifest as START-dependent promotive effects on tumor suppressor activity.

The presence of a START domain suggests DLC proteins may be regulated by a lipophilic ligand, rendering them more easily targetable by natural or synthetic small molecules. However, while there is considerable *in vivo* and *in vitro* evidence for them as bonafide tumor suppressors, the role of their START domain remains poorly understood. In fact, the residues mediating START function remain virtually unknown. This is a key bottleneck for targeted modulation of DLC tumor suppressive activity. Here we use an evolutionary approach to identify conserved residues in DLC START domains that are also mutated in the Catalogue of Somatic Mutations in Cancer (COSMIC) dataset. We find that conserved residues of DLC-1 and DLC-2 START domains are highly-overrepresented in tumor samples from a broad spectrum of cancers. Moreover, using comparative structural modeling, we identify a set of three residues, present in both DLC-1 and DLC-2 START domains, that cluster and interact at the tertiary level. Mutations identified in COSMIC are predicted to break these non-covalent interactions, and disrupt START tertiary structure and function. Our results support the notion that the START domain is involved in DLC-mediated tumor suppression, and that DLC proteins may thus be particularly tractable therapeutic targets. Importantly, these residues would not have emerged from traditional scans for mutational hotspots. As such, we propose that tandem evolutionary and structural analyses could help unmask hidden cancer-relevant mutations within the COSMIC dataset.

## 2. Results and Discussion

### 2.1 Mutations in conserved residues of DLC-1 and DLC-2 START domains are overrepresented in tumors

COSMIC contains numerous missense, nonsense, and frameshift mutations in the START domains of DLC-1, DLC-2, and DLC-3 [36]. We focused on missense mutations, as the latter two are more likely to yield strong effects that confound specific contributions of the DLC START domain. We found 47, 33, and 54 missense mutations falling within the DLC-1, DLC-2, and DLC-3 START domains, respectively (**Table S1-S3**). These mutations do not cluster into hotspots, as a Kolmogorov-Smirnov test for uniformity failed to reject the null hypothesis (**Table S4**). Although uniformly distributed, we hypothesized that they may nonetheless be disproportionately affecting functional residues.

To identify functional residues of DLC START domains, we performed a multiple sequence alignment (MSA), as evolutionary conservation is a strong indicator of critical residues within protein domains [37]. START domains were selected from 123 DLC-1, DLC-2, and DLC-3 orthologs from 46 vertebrate species spanning 450 million years of divergent evolution. Using a stringent 98% threshold, we identified 41 residues (~20%) within the 206 amino acid DLC START domains that are identical in ≥98% of sequences (**Figure 2;** see **Figure S1** for full species MSA). Missense mutations from COSMIC were placed onto the MSA to score whether they preferentially affected these conserved residues (**Figure 3A**). DLC-1 had 17 mutations falling in 11 conserved residues, DLC-2 had 13 mutations falling in 10 conserved residues, and DLC-3 had 5 mutations falling in 5 conserved residues (**Figure 3; Tables S1-S3**). This corresponds to 36.2%, 39.3%, and 9.3% of total mutations in DLC-1, DLC-2, and DLC-3 START domains, respectively (**Table S5**). We next tested whether these frequencies deviate from the approximately 20% likelihood of mutations occurring in conserved residues by chance (**Table S5**). Indeed, mutations are significantly overrepresented in conserved residues of both DLC-1 (p-value = 0.006) and DLC-2 (p-value = 0.005) START domains (**Figure 3B; Tables S5-S7**). By contrast, mutations of the DLC-3 START domain are not overrepresented in conserved residues (p-value = 0.052). These findings are consistent with the frequent downregulation of DLC-1 and DLC-2, but not DLC-3, in many cancers [12]. Taken together, these evolutionary analyses identify conserved residues in DLC-1 and DLC-2 START domains which may be linked to cancer phenotypes.

**Figure 2.**
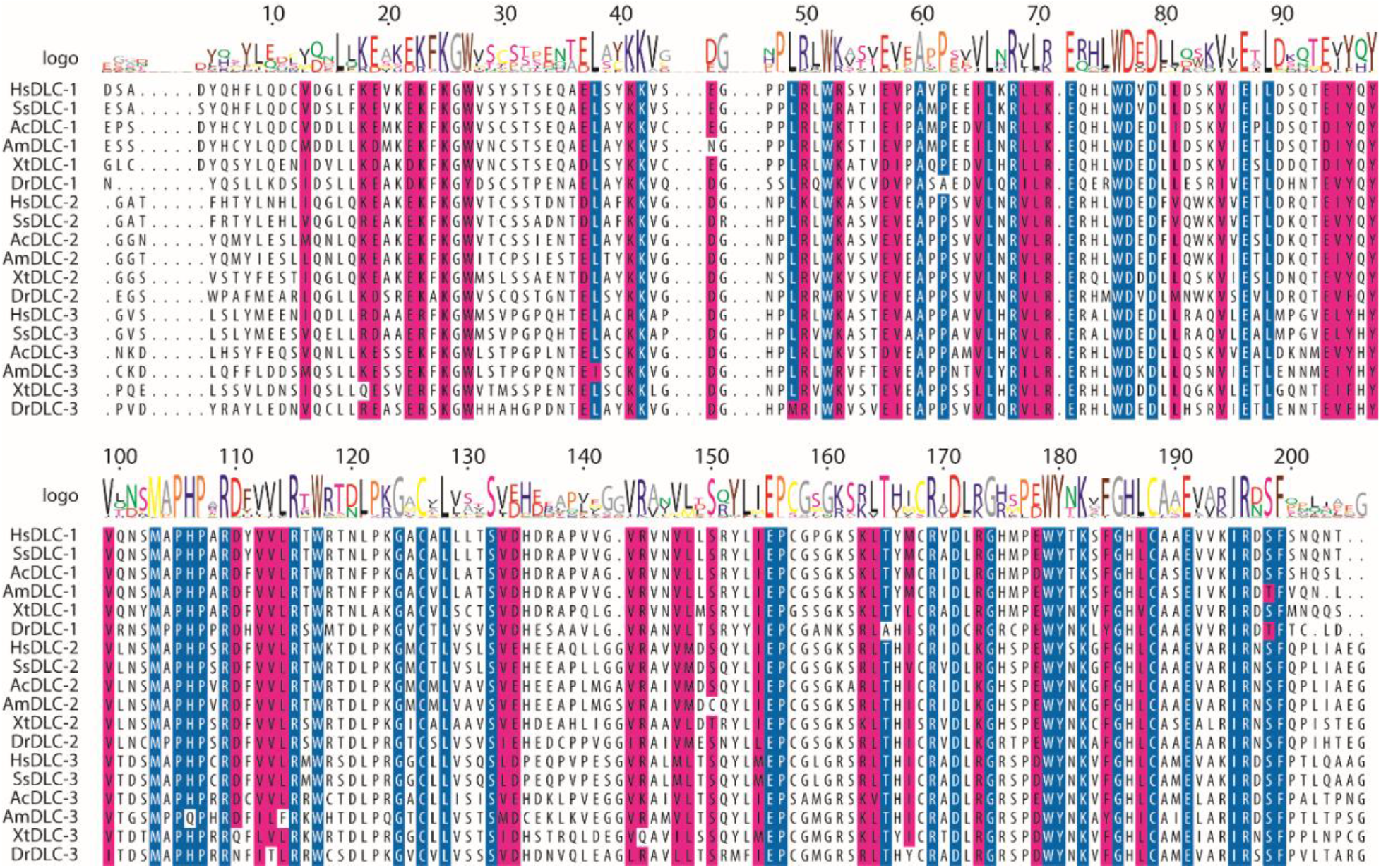
Representative subset of the multiple sequence alignment (MSA) of START domains from DLC-1, DLC-2, and DLC-3 orthologs. Species include Homo sapiens (Hs), Sus scrofa (Ss), Aquila chrysaetos (Ac), Alligator mississippiensis (Am), Xenopus tropicalis (Xt), and Danio rerio (Dr). Blue indicates residues that are identical in ≥98% of sequences. Magenta indicates residues with similar physiochemical properties in ≥98% of sequences. White represents non-conserved residues. Logo represents the consensus residue(s) of the 123 sequences used in the full alignment (see **Figure S1**).

**Figure 3.**
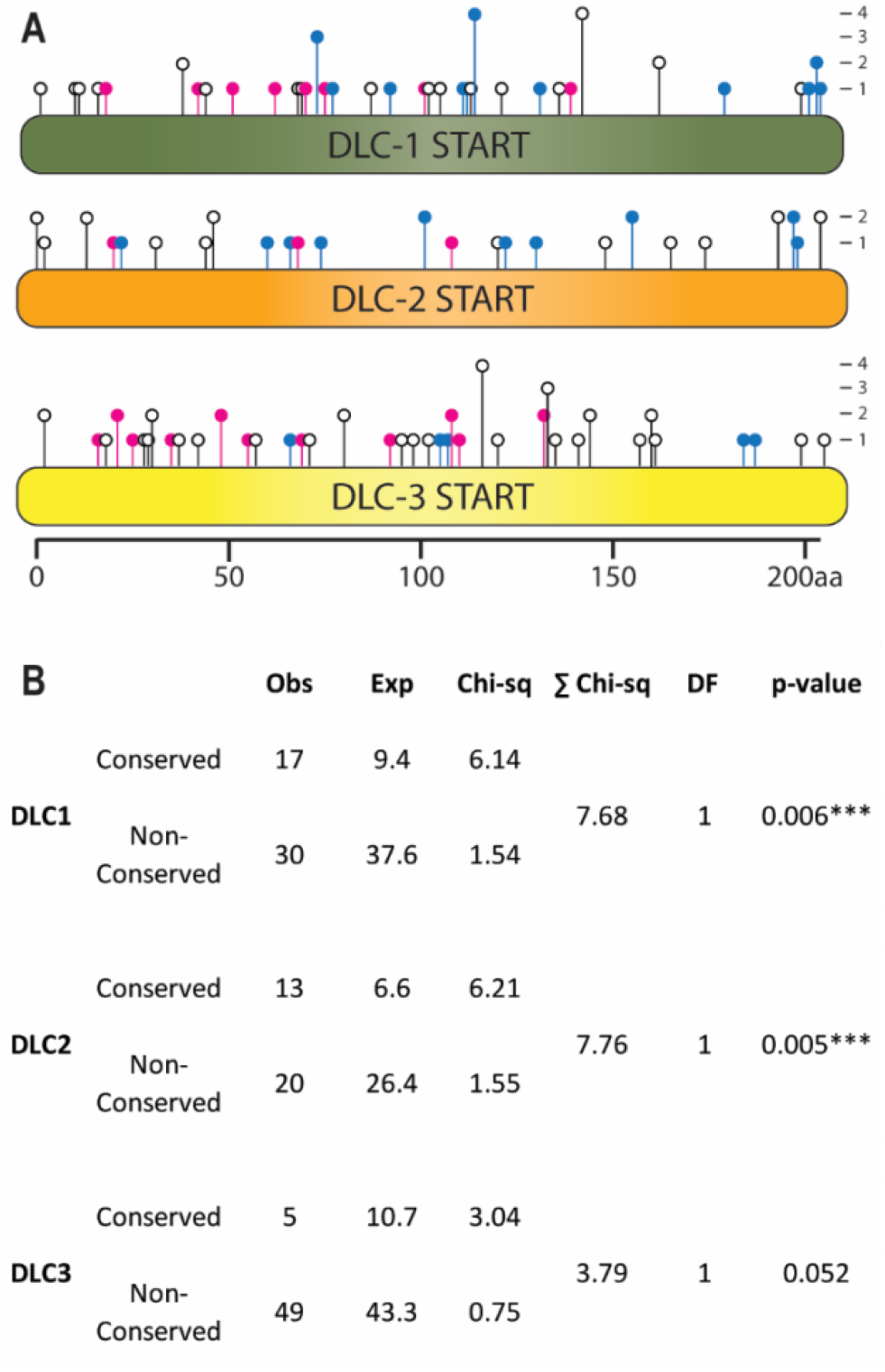
COSMIC missense mutations localizing to conserved residues of DLC-1, DLC-2, and DLC-3 START domains. (**A)** Lollipop plot displaying the location and frequency of missense mutations. Mutations occur in identically conserved (blue), physiochemically conserved (magenta), or non-conserved residues (white). (**B)** COSMIC missense mutations are overrepresented in the evolutionary-conserved residues of the START domains of DLC-1 and DLC-2, but not DLC-3. *** *P* < .001, chi-square test.

### 2.2 Conserved residues form multiple bonds within the START tertiary structure

Conservation of residues implies critical roles in function [37]. For example, an arginine residue near the opening of the START cavity is highly-conserved and known to be required for START activity [38]; this residue also accrued mutations in multiple independent patient samples (DLC-1 R988; **Table S1**). In addition, mutation of the conserved E966 residue in DLC-1 was recently shown to impair tumor suppressor activity [15] (**Table S1**). The recovery of multiple residues with known roles in either cancer or START activity supports the biological relevance of the residues identified by our analyses. Most conserved DLC START residues are uncharacterized however, and it is formally possible that amino acid substitutions have only minimal effects on tertiary structure [39]. We thus sought to predict their impact on structure and function of DLC START domains using comparative structural modeling. As mutations in the DLC-3 START domain were not overrepresented in COSMIC, it was excluded from these structural analyses.

Mutations in COSMIC falling within conserved residues were mapped to homology models of the DLC-1 and DLC-2 START domains (**Figure 4**). Strikingly, three conserved residues with missense mutations in COSMIC colocalized to the same positions within the DLC-1 and DLC-2 START domains: R947, S1077, and F1078 of DLC-1 (**Figure 4A,B**), and R969, S1100, and F1101 of DLC-2 (**Figure 4C,D**). The conserved R947/R969 residues are part of helix-2, and are in close proximity to the S1077/S1100 and F1078/F1101 residues, which sit near the end of the C-terminal helix-3. Given the degree of conservation and close proximity in both DLC-1 and DLC-2, we focused our subsequent analyses on these arginine, serine, and phenylalanine residues.

**Figure 4.**
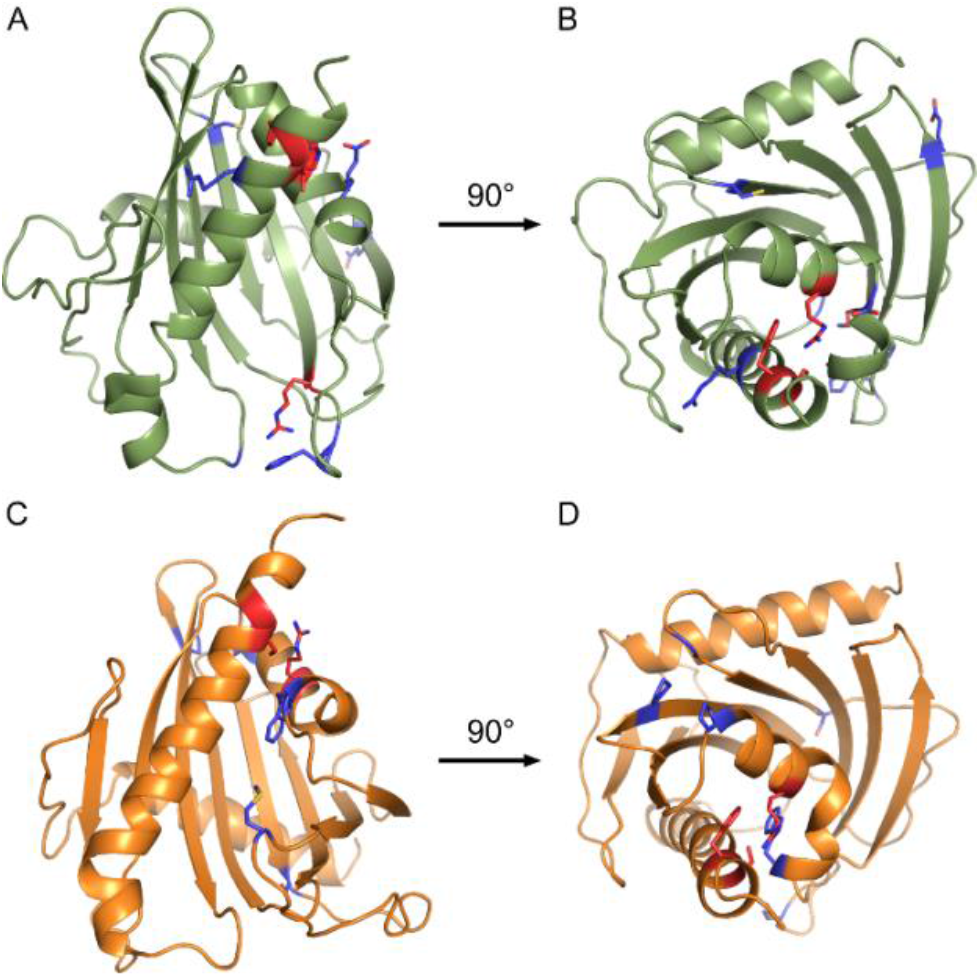
Homology models of DLC-1 and DLC-2 START domains. (**A,C**) Side and (**B,D**) top down views of (**A,B**) DLC-1 and (**C,D**) DLC-2 START domains. Blue indicates conserved residues with missense mutations in cancers. Red indicates conserved arginine, serine, and phenylalanine residues mutated in both DLC-1 and DLC-2, as well as the highly-conserved arginine in DLC-1 (R988) mutated in numerous cancers.

The close proximity of these residues in three-dimensional space presents the possibility that they may interact at the tertiary level. We examined this idea using Crystallographic Object-Oriented Toolkit (CooT), and identified multiple predicted interactions between these residues, as well as between these residues and their neighbors. For instance, the R947 residue of DLC- 1 forms a cation-π interaction with the aromatic ring of F1078 (3.3 Å; **Figure 5A-C**). Cation-π interactions in proteins generally occur between a positively charged amino acid and the π face of an aromatic ring, and are key contributors to protein structure [40, 41]. In addition to this cation-π interaction, R947 also forms a hydrogen bond with Q1081 (3.1 Å). Finally, S1077 does not directly interact with either R947 or F1078, but instead forms a hydrogen bond with K1073 within helix-3. In the case of DLC-2, R969 and F1101 in DLC-2 similarly form a cation-π interaction, although the strength of this interaction is predicted to be weaker (3.8 Å; **Figure 6A-C**). In contrast to S1077 of DLC-1, S1100 in DLC-2 interacts with R969 via hydrogen bonding at both its carboxy backbone (2.9 Å) and side chain (3.2 Å) of S1100. Thus, while not apparent from the primary sequence, three-dimensional homology models reveal these residues appear to function together at the tertiary level.

**Figure 5.**
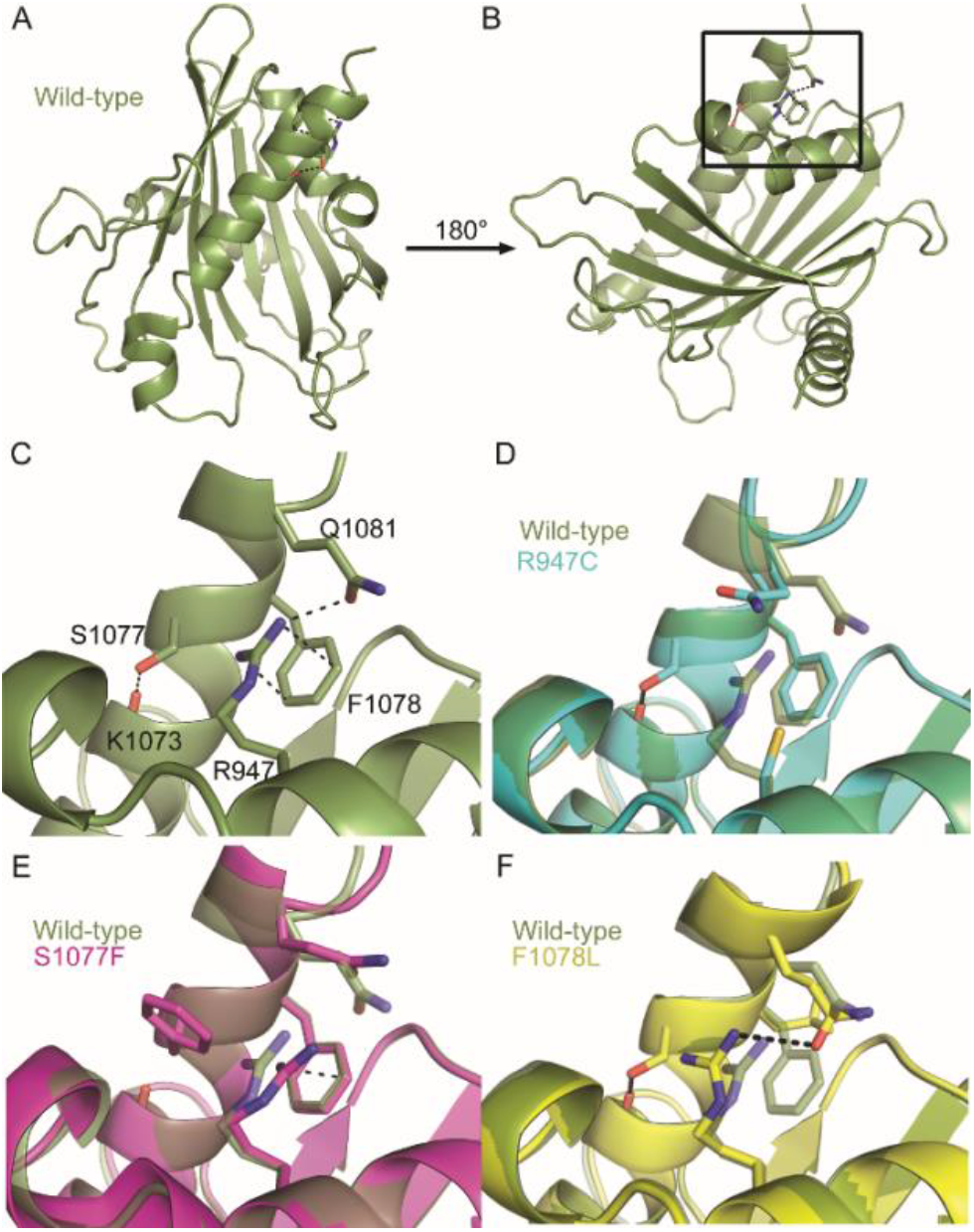
Consequences of COSMIC missense mutations on the tertiary structure of the DLC-1 START domain. (**A-C)** Conserved residues colocalizing near the C-terminal helix form multiple tertiary-level interactions. (**D**) R947C mutation disrupts the cation-π interaction of F1078 and hydrogen bond of Q1081. (**E**) F1078L disrupts the R947 cation-π interaction, but retains the hydrogen bond between R947 and Q1081. (**F**) S1077F retains the cation-π interaction, and disrupts all other bonds. Dotted lines indicate hydrogen bonding and cation-π interactions.

**Figure 6.**
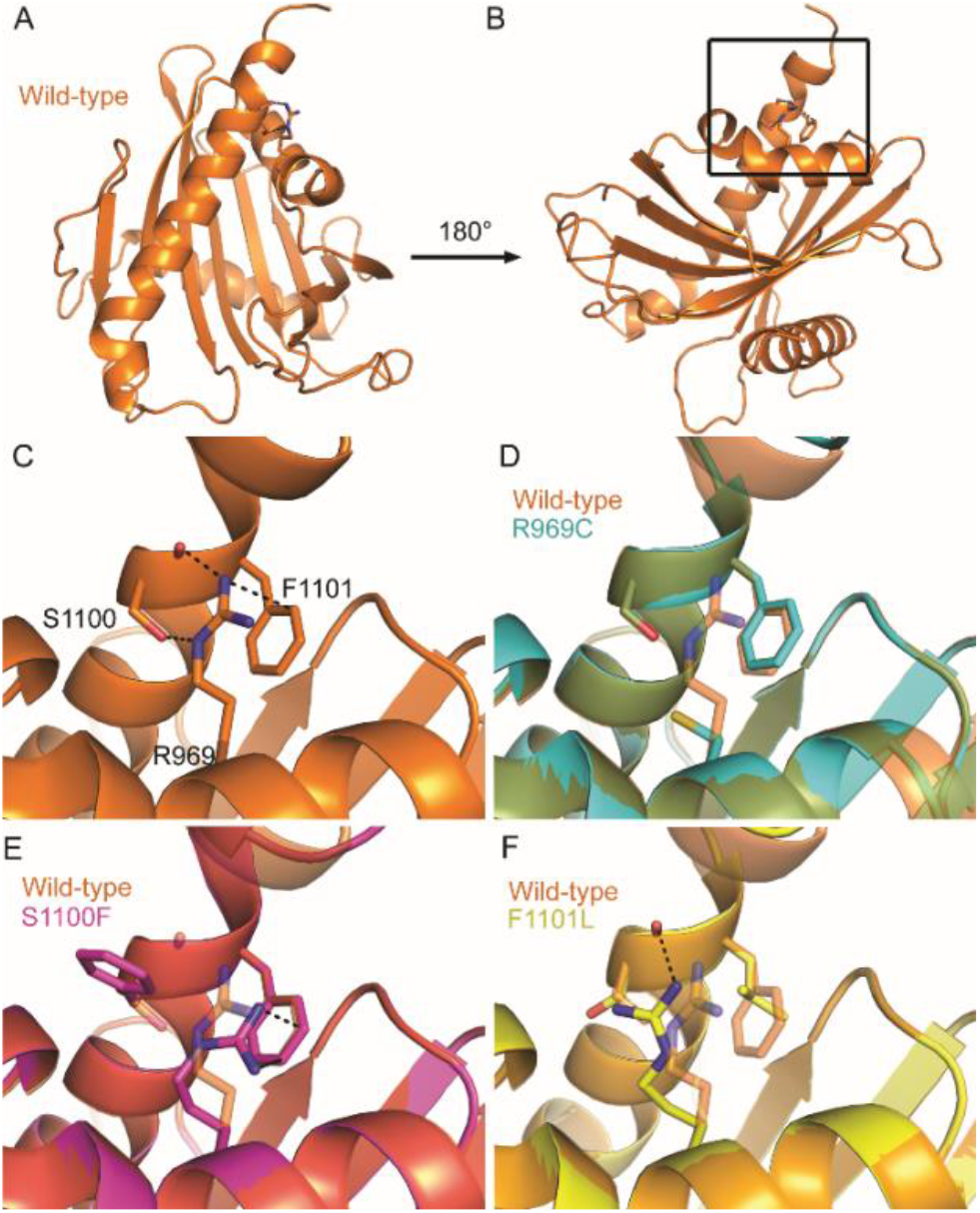
Consequences of COSMIC missense mutations on the tertiary structure of the DLC-2 START domain. **A-C** Conserved residues colocalizing near the C-terminal helix form multiple tertiary-level interactions. (**D**) R969C mutation disrupts all interactions formed between these residues. **E**) F1101L disrupts the cation-π interaction of R969 and the hydrogen bond with the side chain of S1100. Dotted lines indicate hydrogen bonding and cation-π interactions.

### 2.3 COSMIC missense mutations disrupt tertiary-level interactions in DLC-1 and DLC-2 START domains

Their conservation, proximity, and extensive intermolecular interactions suggest that missense mutants in the arginine, serine, or phenylalanine residues would disrupt START structure and function. To test this, we separately replaced each residue with one of the missense mutations identified in COSMIC (**Table S1-S3)**. We then generated new homology models for DLC-1 and DLC-2 START domains and used CooT to determine the likely effect of these mutations on the previously identified noncovalent network.

In DLC-1, the R947C mutation abolishes its cation-π interaction with F1078 and hydrogen bond with Q1081, but leaves the S1077-K1073 hydrogen bond intact (**Figure 5D**). S1077F retains the R947 and F1078 cation-π interaction, but abolishes the S1077F-K1073 hydrogen bond, as well as the hydrogen bond between R947 and Q1081 (**Figure 5E**). Finally, F1078L disrupts its cation-π interaction with R947, but retains the hydrogen bond between R947 and Q1081, as well as the S1077-K1073 hydrogen bond (**Figure 5F**).

In the case of DLC-2, the R969C mutation similarly abolishes its cation-π interaction with F1101, as well as its hydrogen bond to S1100 (**Figure 6D**). S1100F retains the cation-π interaction with R969 and F1101, while abolishing all hydrogen bonds formed with R969 (**Figure 6E**). Finally, F1101L abolishes its cation-π interaction with R969 and disrupts the hydrogen bond between R969 and the side chain of S1100, but retains the hydrogen bond between R969 and the carboxyl backbone of S1100 (**Figure 6F**).

The arginine-to-cysteine mutations in both DLC-1 (R947C) and DLC-2 (R969C) disrupt cation-π interactions, as well as multiple hydrogen bonds. Thus, we predict this substitution to be the most destabilizing in both START domains. This is also in line with global analyses of the COSMIC dataset, which found that arginine-to-glutamine and arginine-to-cysteine substitutions occur with high frequency in driver mutations [42]. The serine-to-phenylalanine mutations (S1077F in DLC-1 and S1100F in DLC-2) also abolish multiple hydrogen bonds and replace a small polar side chain with a large hydrophobic side chain. Thus, this substitution could conceivably destabilize START domains to a similar extent as the arginine-to-cysteine mutations. By contrast, the phenylalanine-to-leucine mutations (F1078L and F1101L) are predicted to be the least destabilizing, as several hydrogen bonds are left intact and both amino acids have similar physicochemical properties [39]. These structural analyses, focusing on a subset of mutations overrepresented in patient tumors, support a role for the START domain in regulating the tumor suppressor activity of DLC-1 and DLC-2.

## 3. Conclusions

Functional residues mediating the tumor suppressive activity of the DLC START domain remain virtually unknown. Here, we undertook a pan-cancer analysis and identified residues within DLC-1 and DLC-2 START domains likely to affect the function of these genes. We also show that COSMIC missense mutations in DLC-1 and DLC-2 START domains are significantly overrepresented in evolutionarily-conserved residues predicted to be critical for proper folding, interaction, and/or activity [37, 38]. Importantly, the relevance of these residues to START and DLC tumor suppressive activities is supported by multiple previous findings [15, 20]. Our analyses thus provide an evolutionary rationale to begin drug targeting of key residues within the conserved START domains of these genes.

Three of these substitutions, occurring in both DLC-1 and DLC-2, disrupt multiple tertiary-level interactions, and are likely to impact START structure. The remaining overrepresented COSMIC missense mutations may similarly impact START activity, or affect other properties such as ligand binding or conformational change. A potential DLC START regulatory mechanism is suggested by studies of both minimal proteins and SMPs, demonstrating that START domains can stimulate regulatory outputs in an allosteric manner [32, 35]. If the DLC START domain functions similarly, it could promote RhoGAP tumor suppressor activity, possibly via allosteric regulation of intra- or inter-molecular interactions.

These data also suggest mutation hotspots might not capture the full spectrum of cancer-relevant mutations in COSMIC. For instance, the conserved arginine, serine, and phenylalanine residues in DLC-1 and DLC-2 appear to work together to stabilize START structure (**Figures 5** and **6**). Single substitutions in any of these residues would have the same structural and functional consequences for DLC START activity, thereby diluting its signal in COSMIC. We propose that, in addition to traditional analyses of mutation frequency, COSMIC should be queried using coupled evolutionary and structure-function analyses. These tandem approaches may help unmask hidden cancer-relevant mutations.

DLC-1 and DLC-2 are frequently down-regulated, as opposed to mutated, in cancer [12, 43]. For instance, DLC-1 expression is reduced 10-fold in lung adenocarcinoma, 3-fold in hepatocellular carcinoma, 4-fold in breast cancer, and 3-fold in colon cancer [12]. Stimulating the activity of residual DLC-1 and DLC-2 proteins through an agonistic START ligand could be a means to improve patient outcomes, particularly if used in combination with current therapeutic approaches. Importantly, agonistic START ligands have already been developed. In plants, signaling of the hormone abscisic acid (ABA) is mediated by its minimal START domain receptor [44]. Opabactin, a rationally designed agonistic START ligand, has a 7-fold stronger affinity than the native ligand, and increases ABA signaling outputs *in vivo* by a factor of 10 [45]. An agonistic ligand designed for the DLC START domain could compensate for reduced outputs, and would be relevant for a number of cancer types. Taken together, our data identify several conserved residues likely to underlie the START-directed regulation of DLC-1 and DLC- 2 tumor suppressive capability. DLC-1 and DLC-2 may thus be high-priority candidates for development of novel therapeutics targeting their START domain.

## 4. Materials and methods

### 4.1 Kolmogorov-Smirnov test of uniformity

Mutation data was obtained from the COSMIC database (v91 GRCh38) for the dominant isoforms of DLC-1 (DLC1_ENST00000358919.6), DLC-2 (ENST00000336934.9), and DLC-3 (ENST00000374599.7). DLC START domains were identified using HHpred [46]: DLC-1 (880 - 1083aa), DLC-2 (903aa-1108aa), and DLC-3 (893aa – 1098aa). Mutations outside of the START domain, as well as nonsense, frameshift, and intronic mutations within the START domain, were excluded from our analyses. To ensure only independent mutation events were measured, duplicate samples from the same patient were removed before analysis. Missense mutations were counted for individual residues, and a one-sample nonparametric test was done using the Kolmogorov-Smirnov test in IBM SPSS Statistics 26. The observed distribution of missense mutations was tested against the null hypothesis that mutations are uniformly distributed along the sample data. Test option settings were set to a significance value of 0.05, a confidence value of 95%, and to exclude cases test-by-test. User-Missing Values were set to “Exclude”.

### 4.2 Multiple Sequence Alignment (MSA)

To build an MSA and identify conserved residues in DLC START domains, orthologs of DLC-1, DLC-2, and DLC-3 were identified using NCBI’s Blastp program against the reference proteins (refseq_protein) database [47].In total, 123 full length sequences were found from 46 vertebrate species. Sequences were aligned using the ClustalW algorithm [48] in the MEGA X software [49], and trimmed to exclude residues falling outside the START domain. Conservation of residues was analyzed using R and “msa” package from Bioconductor [50]. A stringent 98% consensus threshold was used, allowing for only 2 of the 123 sequences to contain a non-similar residue at a given position. This threshold was selected to account for rare lineage specific mutations that may be permissive to changes in critical residues, but not representative of most sequences [51]. Residues were then shaded according to similarity and a consensus logo was added using R and the “msa” package from Bioconductor. Finally, this file was exported in .tex format and compiled using TeXworks (**Figure S1**).

### 4.3 Mutation counts and chi-square analysis

The expected number of mutations in conserved residues assuming a random distribution was calculated by using the percentage of identical residues multiplied by the total number of mutations in START domain of each paralog.

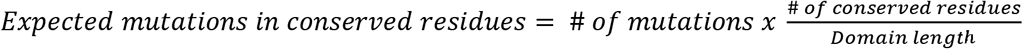

The expected number of mutations in non-conserved residues was determined by subtracting the expected number of mutations in conserved residues from the total number of mutations. Chi-square values were calculated using the formula:

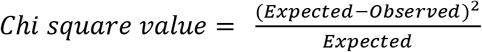

These summed chi-square values were then calculated using the website: https://www.socscistatistics.com/pvalues/chidistribution.aspx

### 4.4 Homology modeling and identification of tertiary-level interactions

Homology modeling of the each START domain was performed using the Max Planck Institute Bioinformatics Toolkit [46]. START domain sequences of DLC-1 and DLC-2 were separately queried using HHpred against the Protein Data Bank (PDB_mmCIF70_20_May) structural database to identify related protein structures for homology modelling. The top 10 structures were identical for both START domains, and included the crystal structure of DLC-2 (2PSO). These 10 protein structures (chain A of 2PSO, chain A 2R55, chain A of 2MOU, chain A of 5I9J, chain A of 6L1M, chain A of 2E3P, chain C of 3P0L, chain A of 1LN1, chain B of 3FO5, chain B of 3QSZ) were forwarded to HHpred-TemplateSelection to generate the MSA. Template alignment was subsequently forwarded to MODELLER to predict the tertiary structures of the DLC-1 and DLC-2 START domains. Structural models were examined using Crystallographic Object-Oriented Toolkit (CooT) to identify predicted interactions formed with the conserved residues using the Environmental Distances tool with a cut-off of 4.0 Å. Mutated DLC-1 and DLC-2 START domain were modeled using the same template alignment as their respective wildtype models, and analyzed for predicted interactions in CooT using the same cut-off value.

## Supplementary Materials

The following are available online at XXX, Figure S1: Full multiple sequence alignment (MSA) of START domains from DLC-1, DLC-2, and DLC-3 orthologs. Tables S1-S7 in Excel.

## Author Contributions

Conceptualization, ASH, RCP, and AYH; methodology, ASH, RCP, RAB, and AYH; software, ASH and RAB; validation, ASH, RCP, RAB and AYH; formal analysis, ASH, RAB, and AYH; investigation, ASH and RCP; resources, ASH, RCP, and AH; data curation, ASH; writing—original draft preparation, ASH and AYH; writing—review and editing, ASH, RCP, RAB, and AYH; visualization, ASH, RAB, and AYH; supervision, AYH; project administration, AYH; funding acquisition, RCP and AYH. All authors have read and agreed to the published version of the manuscript.

## Acknowledgements

We thank members of the Husbands Lab for insightful comments on the work.

## Supplemental Figure and Table Legends

**Figure S1.** Full multiple sequence alignment (MSA) of START domains from DLC-1, DLC-2, and DLC-3 orthologs. Species are listed down the left side of the MSA. Blue indicates residues that are identical in ≥98% of sequences. Magenta indicates residues with similar physiochemical properties in ≥98% of sequences. White represents non-conserved residues. Logo represents the consensus residue(s) of all 123 sequences.

**Table S1** – COSMIC missense mutations and sequence conservation for DLC-1. Conserved and non-conserved residues in the DLC-1 START domain mutated in COSMIC. Features include: residue number, number of missense mutations, identity of introduced substitution, and level of conservation in MSA.

**Table S2** – COSMIC missense mutations and sequence conservation for DLC-2. Conserved and non-conserved residues in the DLC-2 START domain mutated in COSMIC. Features include: residue number, number of missense mutations, identity of introduced substitution, and level of conservation in MSA.

**Table S3** - COSMIC missense COSMIC missense mutations and sequence conservation for DLC-3. Conserved and non-conserved residues in the DLC-3 START domain mutated in COSMIC. Features include: residue number, number of missense mutations, identity of introduced substitution, and level of conservation in MSA.

**Table S4** - Kolmogorov-Smirnov (K-S) test of uniformity for DLC-1, DLC-2, and DLC-3 START domains. K-S test indicates no deviation from uniformity for all DLC START domains. *P* > 0.05.

**Table S5** – Summary of DLC-1, DLC-2, and DLC-3 mutations in conserved and non-conserved residues. Percentage of residues ?98% conserved in all DLC START domains (green), percentage of conserved residues with at least one mutation (pink), percentage of all mutations falling in conserved residues (grey).

**Table S6** – Chi-square input value determination. Calculation of Expected numbers of mutations falling in conserved or non-conserved residues (for chi-square test).

**Table S7** – Chi-square analysis. Chi-square tested demonstrating DLC-1 and DLC-2 START domains accrue significantly more mutations in conserved residues than expected by chance. DLC-3 does not show this effect. \*\*\**P* < 0.001.

